# Multi-omic Profiling of Senescence Following Myocardial Infarction Reveals Dynamic Characteristics in Cardiac Remodeling

**DOI:** 10.1101/2025.08.04.667980

**Authors:** Jielin Deng, Di Ren, Heather Zhou, Yunqiu Jiang, Qiutang Xiong, Yanbo Zhao, Yin Wang, Hao Wu, Makoto Nakanishi, Yingfeng Deng, Zhao V. Wang

**Affiliations:** Department of Diabetes and Cancer Metabolism, Beckman Research Institute, City of Hope National Medical Center, Duarte, California 91010, USA; Irell and Manella Graduate School of Biological Sciences, City of Hope National Medical Center, Duarte, California 91010, USA; Division of Cancer Cell Biology, The Institute of Medical Science, The University of Tokyo, Minato-ku, Tokyo 108-8639, Japan; City of Hope Comprehensive Cancer Center, Duarte, California 91010, USA

## Abstract

Premature senescence is essential for tissue remodeling. Myocardial infarction (MI) induces pathological cardiac remodeling through fibroblast-driven extracellular matrix (ECM) production. The role of senescence in MI-induced remodeling process remains elusive. Here we identify a gradual increment number of senescent cells within the ischemic heart, peaking at day 7 post-MI, in both wild-type and p16^Ink4a^-Cre^ERT2^-mT/mG senescence reporter mice. Lineage tracing shows that senescent cells transition to non-senescent state within 4 weeks after MI. We perform single-nucleus (sn) Multiome and fluorescence-based spatial transcriptomics analyses to profile senescent cells. We next generate a reference (query dataset) based on SPiDER-βGal/p16-EGFP positivity and map it back to the snMultiome dataset. We then deconvolute senescent cells in the integrated dataset using multiple computational algorisms. Through these approaches, we reveal that fibroblasts and its subpopulation-late myofibroblasts (MF)-constitute a major proportion of senescent cells, which functionally reduce ECM production. Importantly, ischemia-induced senescent MF show less soluble collagen production compared to TGF-β1-induced non-senescent MF *in vitro*. At the functional level, depletion of senescent cells *in vivo* augments fibrosis and worsens cardiac myopathy post-MI. Our findings highlight the transient nature of senescent cells in the heart and underscore the importance of dynamic regulation of senescent cells post-MI.

## Main

Persistent senescence is elevated in aging, while premature senescence occurs upon cellular stress and injury beyond aging^1^. Myocardial infarction (MI) is a leading cause of morbidity and mortality worldwide^2^, which triggers various cellular stresses resulting from reduced blood flow and oxygen supply to the heart. This process is accompanied by pathological myocardial remodeling after ischemia-induced cardiomyocyte death and immune cells-triggered inflammation^3^. Several studies have shown that MI causes premature senescence in multiple myocardial cell types, including cardiomyocytes, fibroblasts, and endothelial cells, as reflected by immunofluorescence staining ^4,5^. However, the multicellular dynamics and cellular composition of senescence in this complex pathophysiological context of MI-induced cardiac remodeling remain elusive. Whereas both beneficial and detrimental functions of senescent cells have been reported^6,7^, no systemic and comprehensive studies have been undertaken to elucidate their kinetics, characteristics, and functional implications during cardiac remodeling following MI. Therefore, an essential first step is to precisely identify senescent cells in the infarcted heart. Currently, there is no unique marker to detect senescent cells^8^. Common experimental approaches including SA-β-Gal/SPiDER-β-Gal activity^9,10^ and cell cycle arrest inducer p16^Ink4a^ (encoded by *Cdkn2a*) expression^11,12^ are widely accepted as the most specific markers to characterize senescence. Genetically engineered p16^Ink4a^-related mouse models such as fluorescence-based reporter lines^13,14^ also help to identify cellular senescence *in vivo*.

Spatial transcriptomics integrated with fluorescence staining has been adapted to delineate the characteristics of target cells, which may provide a deeper understanding of senescence in the heart post-MI. Nevertheless, comprehensive mapping of resident cell types from spatial transcriptomics alone remains a challenge. A technical hurdle is that spatial transcriptomics itself lacks the single-cell resolution^15^ and sensitivity to resolve subtle transcriptional differences of fine-grained cell types^16^. It’s therefore essential to utilize computational tools to deconvolute spatial transcriptomics by leveraging cell type signatures from RNA-seq at the single-cell resolution and transferring cell labels into senescent marker fluorescence-labeled spatial transcriptomics data.

Here we conduct time-dependent MI in both wild-type and p16^Ink4a^-Cre^ERT2^-mT/mG senescence reporter mice to investigate the dynamic regulation of senescent cells in the heart in response to MI. We next analyze both single-nucleus Multiome (snMultiome) and fluorescence-labeled (SPiDER-β-Gal staining and p16^Ink4a^-EGFP) spatial transcriptomes, separately or integrated, to deconvolute and explore the features and functions of senescent cells upon MI. Finally, we validate our findings and reveal the role of senescence in MI through both *in vitro* and *in vivo* approaches.

## Results

### Senescent cells transiently accumulate in the infarct region of the heart post-MI

To investigate the dynamic regulation of senescence in the heart in response to MI, left anterior descending (LAD) coronary artery was ligated to induce MI in adult wild-type mice (**Fig. 1a**). At different time points following MI (sham and 3 days, 7 days, 2 weeks, 3 weeks, and 4 weeks post-MI), hearts were harvested for analysis. We found that the highest SA-β-Gal activity was observed within the infarction region on day 7 post-MI compared to other groups (**Fig. 1b**). The SA-β-Gal staining of different cross-sections of the heart on day 7 post-MI was shown in **Extended Data Fig. 1a**, demonstrating that various transverse heart layers exhibit a similar degree of senescence, providing the rationale for the subsequent integrated analysis of snMultiome-seq and spatial transcriptomic-seq data from the same infarcted heart.. In addition, the protein level of several senescent markers, including p16, p21, and p53, within the infarction region began to increase on day 3, reached the peak at day 7 (p16 and p21) or day 14 (p53), and gradually reduced within 4 weeks post-MI (**Fig. 1c**). Moreover, both γH2AX immunofluorescence staining and SPiDER-β-Gal enzymatic conjugates fluorescence staining^17^ showed a significantly higher number of positive cells on day 7 post-MI compared to sham MI (**Fig. 1d and Extended Data Fig. 1b**).

**Fig. 1.**
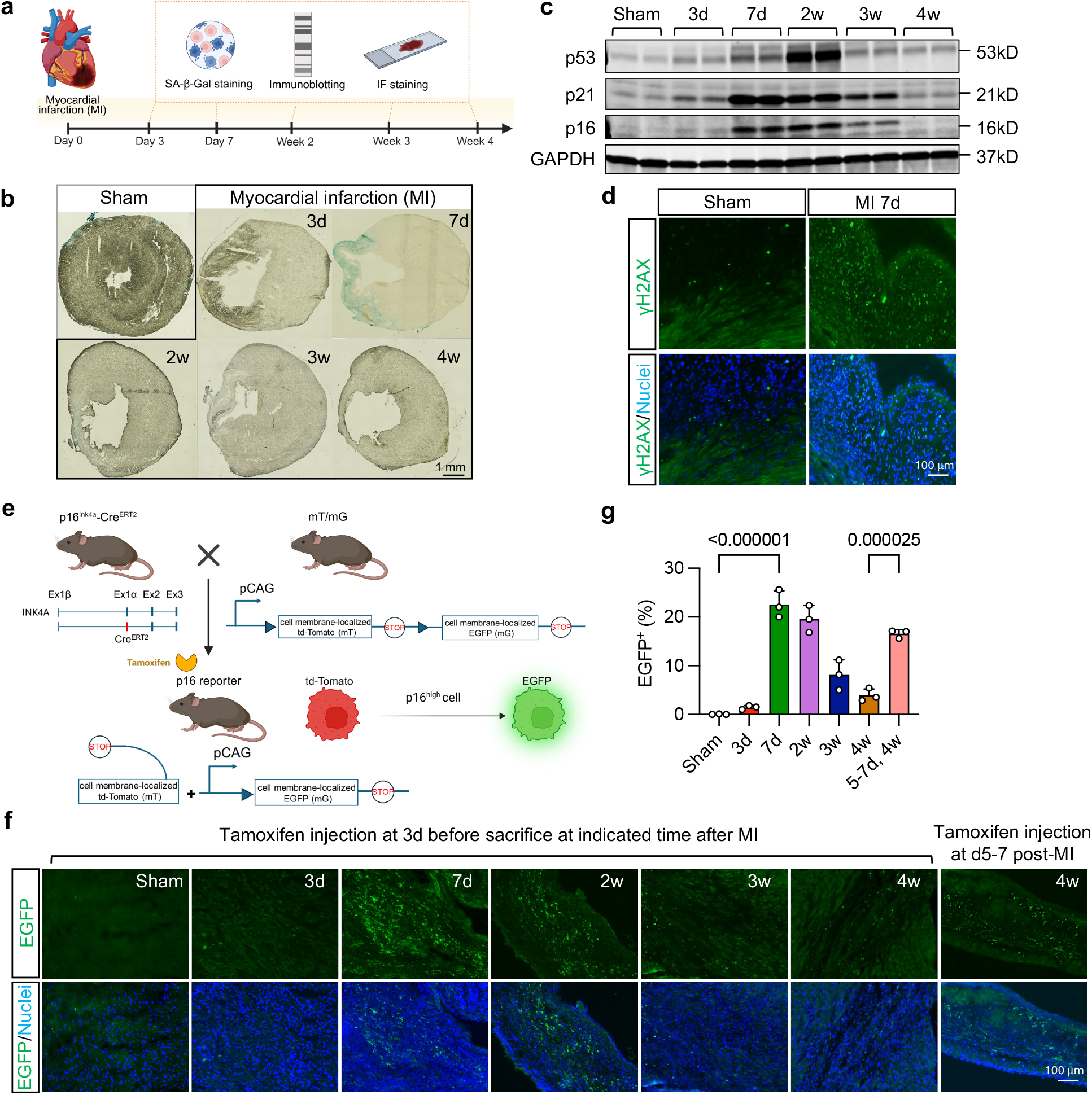
Senescent cells transiently accumulate in the infarct region of the heart post-MI. **a**. Experimental design to evaluate cellular senescence of the heart following myocardial infarction (MI). Senescence-associated β-galactosidase (SA-β-Gal) staining, immunoblotting, and immunofluorescence (IF) staining were performed on days 3 and 7, as well as 2-, 3-, and 4-weeks, post-MI to evaluate cardiac senescence. **b**. SA-β-Gal staining of heart cross-sections at different time points post-MI, showing the highest number of SA-β-Gal positive cells on day 7 post-MI. **c**. Immunoblots of senescence markers p16, p21, and p53 in left ventricular lysates at various time points post-MI. Expression of p16 and p21 peaked at day 7, whereas p53 peaked at day 14, followed by a gradual decline. GAPDH served as a loading control. **d**. Representative γH2AX IF staining of heart sections from sham and day 7 post-MI mice, showing an elevated DNA damage response on day 7 post-MI. **e**. Schematic diagram of the p16 senescence reporter mouse model. p16^Ink4a^-Cre^ERT2^ mice were crossed with mT/mG mice to generate a tamoxifen (TAM)-inducible p16 senescence reporter model, allowing visualization of p16^high^ cells via EGFP fluorescence detection. **f**. Imaging of EGFP positive p16^high^ cells in p16 senescence reporter mice at different time points post-MI. EGFP signals peaked on day 7 post-MI and gradually declined. Lineage tracing of p16^high^ cells revealed stronger EGFP signals at 4 weeks post-MI compared to those observed following TAM injection 3 days prior to the 4 weeks post-MI harvesting time point. **g**. Quantification of EGFP positive cells is shown. One-way ANOVA followed by Tukey post-hoc multiple comparisons test was conducted. Data are represented as mean ± SD.

Next, we took an independent approach to further investigate the dynamic regulation of senescent cells upon MI. We crossed the p16^Ink4a^-Cre^ERT2^ mouse^13^ with the mT/mG mouse^18^ to generate a p16 senescence reporter mouse model, which enables effective labeling of p16^high^ cells with EGFP in the presence of tamoxifen (TAM, 3 consecutive days, daily i.p. injection before harvest) (**Fig. 1e**). We observed a progressive increase of p16^high^ cells over ischemic time, which peaked at day 7, followed by a gradual decrease within 4 weeks (**Fig. 1f and 1g**). Consistently, immunostaining of p16^Ink4a^ of the wild-type mouse heart at different time points following MI showed a kinetic pattern similar to that of EGFP positive cells in p16 reporter mice (**Extended Data Fig. 1c**). In addition, SA-β-Gal staining of the p16 senescence reporter mouse heart confirmed the co-localization of SA-β-Gal positive cells with p16^high^ cells (**Extended Data Fig. 1d and 1e**). These findings demonstrated that p16 and SA-β-Gal are two similar and convincing markers of senescence in the MI mouse model. Finally, to map the fate of senescent cells in the heart post-MI, we conducted a lineage-tracing assay. TAM was administered on days 5-7 to the p16 senescence reporter mouse, and the hearts were harvested at 4 weeks post-MI. We found that p16^high^ cells, which peaked on day 7, were still evidently present 4 weeks post-MI (**Fig. 1f and 1g**). The number of p16^high^ cells were higher than those in mice injected with TAM 3 days prior to the 4 week-harvest, suggesting that most of the p16^high^ cells gradually turn into the non-senescent state. Taken together, these findings indicated that senescent cells in the heart in response to MI manifest dynamic and transient characteristics.

### Fibroblasts are prone to senescence in the heart following MI

To characterize the senescent cells in the heart observed on day 7 post-MI, we set up to profile global cardiac transcriptomics and integrated it with spatial transcriptomics to identify the composition of the senescent cells at the single-cell resolution. To achieve this goal, the heart from day 7 post-MI was divided into two parts below the ligation site. Half of the lower part (remodeling region) was homogenized to generate nuclear suspension for sing-nucleus (sn) Multiome analysis. An infarct heart slice from the upper part was used for CytAssist spatial transcriptomics based on the SPiDER-βGal and p16-EGFP fluorescence in wild-type and p16 senescence reporter mice, respectively (**Fig. 2a**).

**Fig. 2.**
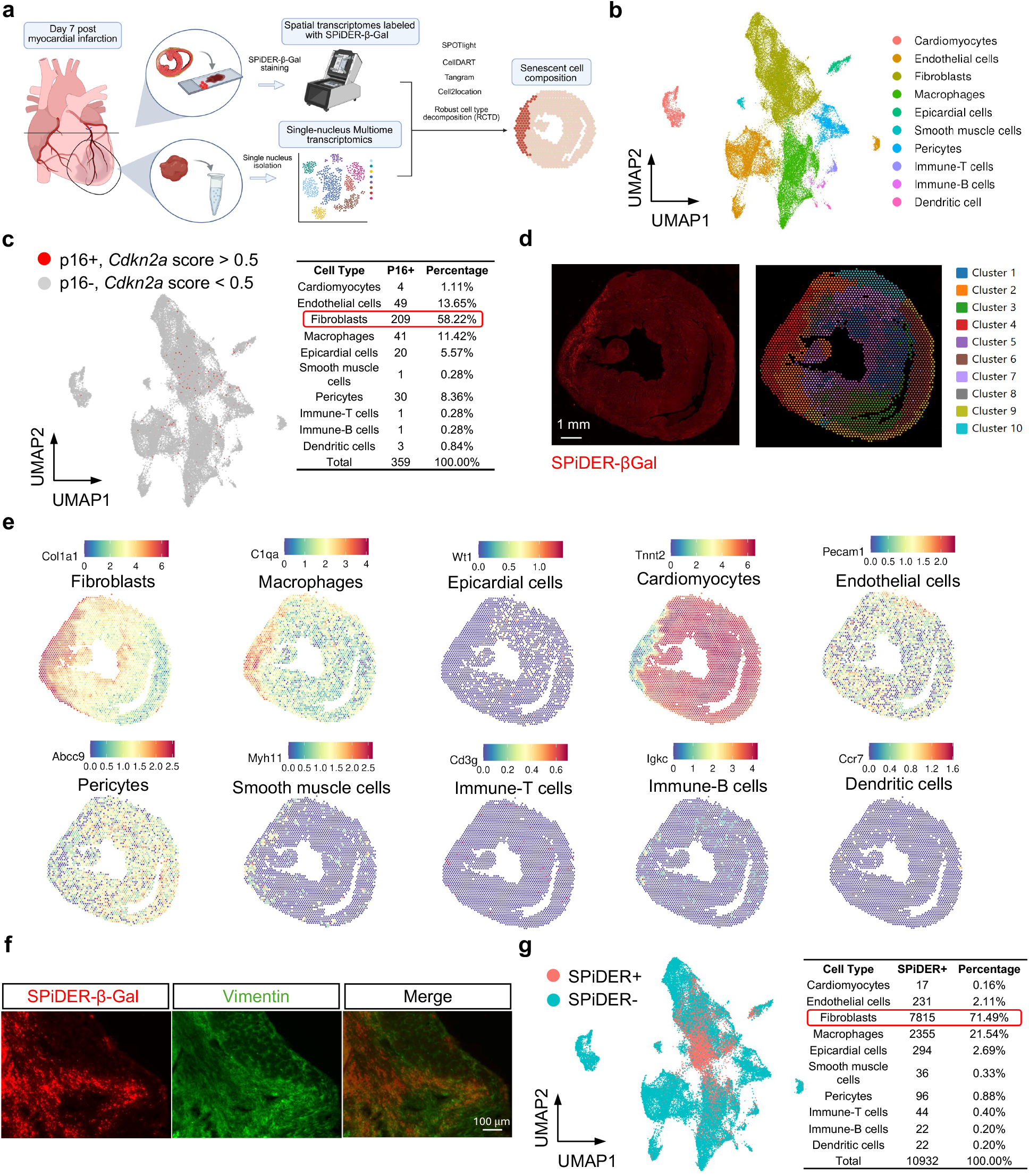
Fibroblasts constitute a major portion of the senescent cell population in the infarcted heart on day 7 post MI. **a**. Schematic diagram of the combinational analysis, integrating spatial and snMultiome transcriptomics datasets. **b**. Single-nucleus (sn) multiome sequencing analysis from the infarcted heart of wild-type and p16 senescence reporter mice day 7 post-MI. WNN identified 10 cardiac cell types. **c**. Distribution of p16 positive senescent fibroblasts (*Cdkn2a* gene filter score > 0.5) across all the 10 cardiac cell types, with fibroblasts comprising 58.22% of the total senescent cells. **d**. Unsupervised spatial transcriptomic analysis of the infarcted heart on day 7 post-MI in the wild-type mouse, identifying 10 clusters. **e**. Fibroblasts (*Col1a1*) were predominantly localized in the ischemic region (cluster 4) where the most SPiDER-βGal positive senescent cells were. **f**. SPiDER-β-Gal (red) colocalized with vimentin (green) in the infarcted heart section from the wild-type mouse on day 7 post-MI, implying that senescent cells are predominantly fibroblasts. **g**. Mapping the SPiDER-βGal fluorescence-based spatial transcriptomics to snMultiome data, showing that fibroblasts comprise 71.49% of the total SPiDER-βGal positive cells.

We first performed snMultiome sequencing, encompassing Assay for Transposase-Accessible Chromatin (snATAC) and Gene Expression (snRNA) sequencing on heart samples from the wild-type mice and p16 senescence reporter mice using the 10x Genomics Chromium platform. Following the quality control procedure, nuclear dataset consisting of a total of 44,573 nucleus was used to generate 31 unsupervised clusters. Weighted nearest neighbor (WNN) procedure was applied to snMultiome RNA-seq data and identified 10 cell types in the heart, including fibroblasts, macrophages, endothelial cells, pericytes, cardiomyocytes, epicardial cells, smooth muscle cells, immune-T cells, immune-B cells, and dendritic cells (**Fig. 2b and Extended Data Fig. 2a**). The expression level of *Cdkn2a* gene (encoding p16) was pulled out as the senescence marker gene in snMultiome RNA-seq data to obtain a single senescence score for each cell by “AddModuleScore”. Cells with the score larger than 0.5 were identified as p16 positive senescent cells, and less than 0.5 were identified as p16 negative non-senescent cells. The proportion of senescent vs. non-senescent cells in each cell cluster revealed that fibroblasts were the predominant cell type of senescent cells, representing 58.22% of the total p16 positive senescent cells (**Fig. 2c**).

We then analyzed the spatial transcriptomics data in an unsupervised manner. We identified 10 clusters in both wild-type and p16 senescence reporter mice (**Fig. 2d and Extended Data Fig. 2b**). Notably, cluster 4 (ischemic region) contained the highest number of SPiDER-βGal positive cells in the wild-type mouse heart, marking an area of cellular senescence (**Fig. 2d**). Fibroblasts (*Col1a1*) and macrophages (*C1qa*) were predominantly localized in clusters 4 and 10, representing the ischemic and border zones, respectively (**Fig. 2e**). In contrast, cardiomyocytes (*Tnnt2*) were present in all clusters except cluster 4. Other cell types were distributed across different clusters, including epicardial cells (*Wt1*), endothelial cells (*Pecam1*), pericytes (*Abcc9*), smooth muscle cells (*Myh11*), T cells (*Cd3g*), B cells (*Igkc*), and dendritic cells (*Ccr7*) (**Fig. 2e**). In the p16 senescence reporter mouse heart, p16^high^ cells were identified by EGFP fluorescence, enabling direct spatial transcriptomics analysis (**Extended Data Fig. 2b**). Similarly, cluster 2 (ischemic zone) was where p16^high^ cells were present. Fibroblasts and macrophages were located predominantly or scattered within ischemic zone (cluster 2) and border zone (cluster 9 and 10) (**Extended Data Fig. 2c**), whereas cardiomyocytes were largely located in remote zone (all clusters except for cluster 2). These spatial transcriptomics data suggested that the senescent cells are primarily composed of fibroblasts and macrophages, with fibroblasts being the most predominant cell type under MI-induced stress. Additionally, we performed fluorescence co-staining of SPiDER-β-Gal with vimentin, a mesenchymal cell marker in wild-type mouse hearts on day 7 post-MI. SPiDER-β-Gal positive cells were primarily colocalized with fibroblasts (**Fig. 2f**). Additionally, co-localization of γH2AX with vimentin was evident in the ischemic zone of the heart (**Extended Data Fig. 2d**).

After snMultiome transcriptomics and spatial transcriptomics analyses of senescent cells, we next aimed to integrate these two types of data and deconvolute senescent cells at a higher resolution. We first mapped a reference dataset derived from the spatial transcriptomics data, categorized based on senescent marker fluorescence (SPiDER-β-Gal/p16 EGFP positivity). We then projected it back to the snMultiome data to establish a new query dataset. The SPiDER-β-Gal-/p16 EGFP positive cells in the query dataset showed that fibroblasts accounted for 71.49% and 55.05% of the fluorescence positive cells, respectively (**Fig. 2g and Extended Data Fig. 2e**).

Next, to further resolve fine-grained cell types in our spatial transcriptomics data, we applied 5 different algorithms, including non-negative matrix factorization (NMF)-based SPOTlight^19^, probabilistic-based RCTD^20^ and Cell2location^16^, and deep learning-based Tangram^21^ and CellDART^22^, to the integrated dataset. We created a comprehensive cellular map of the infarcted heart by deconvolution of the 10 cell types with higher resolution and sensitivity using the snMultiome transcriptomics data as references. Cell2location outputs estimated cell abundances whereas the other 4 methods output estimated cell proportions. SPiDER-β-Gal and EGFP positive spots from the wild-type and p16 senescence reporter mouse hearts (254 spots and 200 spots after quality control, respectively) were selected manually and pulled out to calculate cell proportions of senescent cells (**Extended Data Fig. 3a**). Cell abundances outputted by Cell2location were converted to proportions within spots. The cell type composition in these spots is shown in **Fig. 3a-e** (for the SPiDER-β-Gal-labeled wild-type mouse heart) and **Extended Data Fig. 3b-f** (for the p16 senescence reporter mouse heart). All 5 algorithms revealed the highest proportion of fibroblasts within the fluorescence positive spots in the spatial coordinates of the ischemic region in Visium sections from both wild-type and p16 senescence reporter mouse hearts (**Fig. 3f and Extended Data Fig. 3g**). To evaluate the degree of agreement among the 5 methods, Pearson correlation coefficient (PCC) was calculated for each pair of methods, revealing strong agreement across all methods (**Fig. 3g and Extended Data Fig. 3h**). Taken together, the new query dataset and all the algorithms consistently identified fibroblasts as the predominant senescent cell type in the heart on day 7 post-MI.

**Fig. 3.**
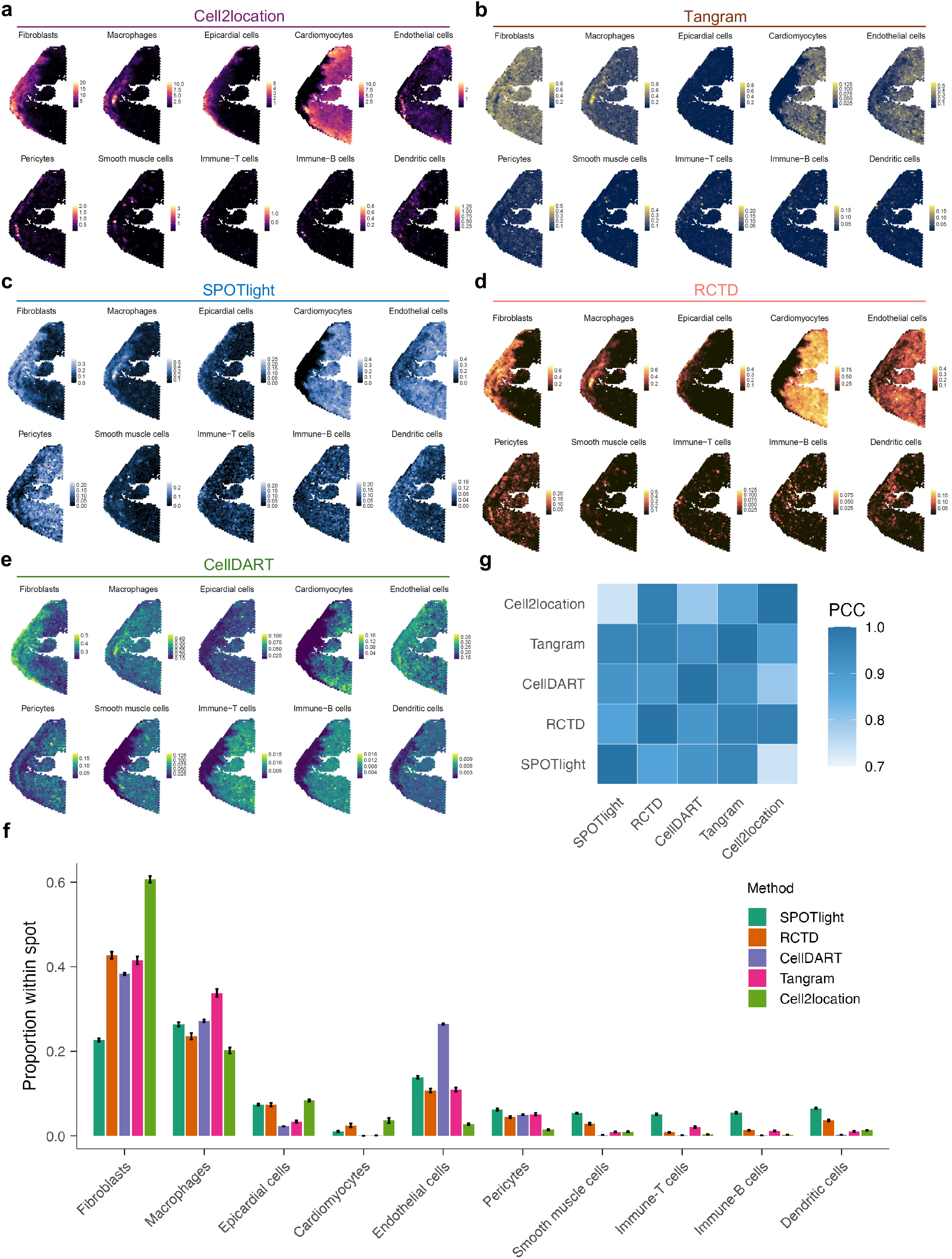
Cellular deconvolution of SPiDER-βGal positive cells confirms fibroblasts as the predominant senescent cell type in the heart post-MI. **a-e**. Cellular deconvolution using Cell2location (**a**), Tangram (**b**), SPOTlight (**c**), RCTD (**d**), and CellDART (**e**) to estimate the proportion of Visium spots in the infarcted heart from the wild-type mouse on day 7 post-MI. **f**. Statistical analysis of cell type compositions predicted by the 5 deconvolution algorithms in SPiDER-β-Gal positive spots confirmed fibroblasts as the predominant senescent cell type. **g**. Pearson correlation coefficient (PCC) was calculated to assess the agreement between each pair of deconvolution algorithms in predicting SPiDER-β-Gal positive senescent cell types in the infarcted wild-type mouse heart. Among the 5 algorithms, RCTD and Tangram showed the highest overall correlation with others. Average PCC values were as follows: SPOTlight, 0.87; RCTD, 0.93; CellDART, 0.89; Tangram, 0.93; Cell2location, 0.85.

### Senescent myofibroblasts manifest reduced collagen production capability upon MI

Different senescent cell types may play distinct roles during MI-induced cardiac remodeling. Since fibroblasts constitute a large proportion of senescent cells, we next focused on senescent fibroblasts within the ischemic region of the infarcted heart to elucidate their roles in pathological cardiac remodeling. Fibroblasts following MI can be categorized into 4 stages^23^: quiescent fibroblasts (quiCF) (day 0), which maintain myocardial tissue function and regulate slow extracellular matrix (ECM) turnover; activated fibroblasts (actCF) (days 1-3), which initiate inflammation and exhibit increased proliferation and migration; myofibroblasts (MF) (days 3-7), derived from actCF, which drive fibrotic tissue formation while maintaining ECM turnover; and matrifibrocytes (MaF) (days 14-28), which contribute to reactive fibrosis and scar maturation. We conducted further sub-clustering analysis and identified 8 distinct fibroblast subtypes across the 4 stages (**Fig. 4a**). Next, we analyzed the distribution and abundance of p16 positive senescent cells (*Cdkn2a* gene filter score > 0.5) within the fibroblast subpopulations. Notably, these p16 positive senescent cells were primarily late MF, comprising 68.72% of the total senescent fibroblasts (**Fig. 4b**). In concordance with this finding, SPiDER-β-Gal/p16 EGFP positive cells in late MF comprised of 71.86% (**Fig. 4c**) and 52.90% (data not shown), respectively, of the total positive cells in the new query snMultiome dataset. These results were further confirmed by the colocalization of γH2AX and the myofibroblast marker α-SMA with immunofluorescence staining (**Fig. 4d**).

**Figure 4.**
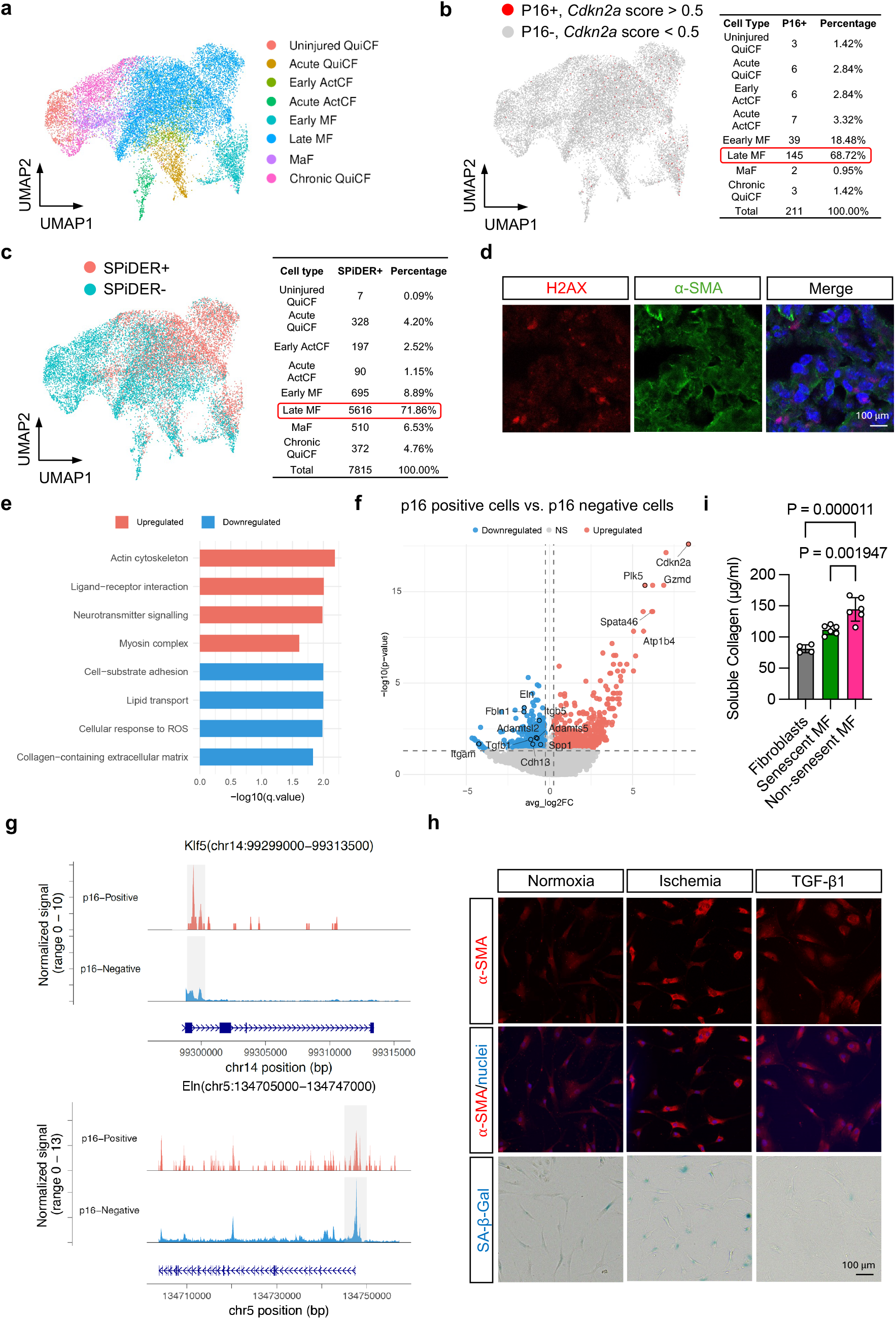
Senescent myofibroblasts exhibit reduced collagen production capability upon MI. **a**. Sub-clustering analysis identified 8 distinct fibroblast subpopulations. **b**. Distribution of p16 positive senescent fibroblasts (*Cdkn2a* gene filter score > 0.5) across fibroblast subpopulations, with late myofibroblasts (MF) comprising 68.72% of the total senescent fibroblasts. **c**. Projection of the SPiDER-β-Gal-labeled spatial transcriptomics dataset to the snMultiome transcriptomics dataset, showing that the late MF accounted for 71.86% of all the SPiDER-β-Gal positive cells in the query dataset. **d**. γH2AX (red) colocalized with α-SMA (green) from the infarcted wild-type mouse heart on day 7 post-MI, confirming that the MF subpopulation constitutes the large proportion of senescent fibroblasts. **e**. Enrichment analysis of differentially expressed genes (DEGs) between p16 positive and p16 negative late MF. **f**. Volcano plot showed the downregulated fibrogenic genes in senescent late MF, indicating a reduced fibrogenic phenotype. **g**. Differentially expressed peaks (DEPs) showed that increased chromatin accessibility of the anti-fibrotic transcription factor (TF) KLF5 and decreased chromatin accessibility of fibrotic gene Eln at the transcription start site in senescent late MF compared to non-senescent late MF. **h**. *In vitro* simulated ischemia increased SA-β-Gal positive cells in human cardiac fibroblasts (HCF) compared to normoxia, whereas TGF-β1 treatment showed no significant effect. These senescent HCF showed enhanced α-SMA positive immunofluorescence signal compared to TGF-β1-induced non-senescent MF. **i**. Statistical analysis showed reduced soluble collagen production in simulated ischemia-induced senescent MF compared to TGF-β1-induced non-senescent MF. One-way ANOVA followed by Tukey post-hoc multiple comparisons test was conducted. Data are represented as mean ± SD.

Transient fibroblast senescence has been suggested to play a short-term role in modulating the fibrotic response during wound healing^24,25^. To explore the role of senescence of late MF in the heart post-MI, differentially expressed genes (DEGs) between p16 positive and p16 negative late MF were applied to pathway enrichment analysis. We found that the DEGs were mainly enriched in collagen-containing ECM, cellular response to reactive oxygen species, and actin binding (**Fig. 4e**). Among the downregulated genes in senescent late MF, multiple fibrogenesis genes were identified (**Fig. 4f**). Next, transcription factor (TF) motif enrichment analysis of differentially expressed peaks (DEPs) from snMultiome ATAC-seq data revealed increased chromatin accessibility of KLF5 (ranking 6, by adjust *p* value), a well-known anti-fibrotic TF, in senescent late MF (data not shown). DEPs data showed increased peaks of *Klf5* and decreased peaks of downstream fibrotic gene *Eln* in senescent late MF compared to non-senescent late MF (**Fig. 4g**). Taken together, these findings suggest that senescent late MF may be implicated in downregulating collagen production.

To validate these findings experimentally, we performed simulated ischemia to induce senescence in human cardiac fibroblasts (HCF). A significant increase in SA-β-Gal positive cells compared to the normoxia group was observed (**Fig. 4h**). In contrast, MF differentiation induced by TGF-β1 did not result in a noticeable difference in SA-β-Gal positivity (**Fig. 4h**). As expected, induction of HCF senescence generated α-SMA positive, contractile cells, with molecular and ultrastructural MF features (**Fig. 4h**), similar to TGF-β1 treatment^26^. However, soluble collagen assay revealed that ischemia-induced senescent MF exhibited reduced collagen deposition compared to TGF-β1-induced non-senescent MF (**Fig. 4h**). Moreover, daily treatment with a senolytics ABT263 (days 4-7 post-MI) in wild-type mice reduced SA-β-Gal positive senescent cells in the heart on day 7 post-MI (**Extended Data Fig. 4a**), confirming its senescent cell-eliminating effects. Daily treatment of ABT263 (days 4-10 post-MI) led to increased fibrosis (**Extended Data Fig. 4b**), and worsened cardiac function (**Extended Data Fig. 4c**) 3 months post-MI. Collectively, these findings suggest that senescent late MF may exhibit a resolved fibrogenic phenotype and reduce ECM deposition, thereby limiting excessive fibrosis and preserving cardiac function during cardiac remodeling after MI.

## Discussion

Our results demonstrated that during cardiac remodeling upon MI, senescent cells transiently peak in the ischemic region on day 7 post-MI, and most of them transition to the non-senescent state over time. We showed that several cell types exhibit senescent characteristics, with fibroblasts and its subpopulation late MF being the most abundant one, as identified through multimodal analysis. Importantly, senescent late MF exhibit reduced collagen production capability, which may limit excessive fibrosis during cardiac remodeling. Taken together, these results highlighted an unexplored role for a stress-induced transient senescence-like state in regulating cardiac remodeling following MI.

Premature senescence is widely considered an irreversible, growth-arrest state^27^. However, recent studies suggested that the full-featured senescence is not necessarily a permanent endpoint but potentially transient, depending on essential maintenance components^28^. Here we showed transient senescence in the infarcted mouse heart over ischemic time. Understanding the endpoint of this transient nature is crucial for gaining insight into the functional outcomes of senescent cells during the remodeling process. Our lineage tracing study demonstrated that most p16^high^ senescent cells from day 7 post-MI were still abundant at 4 weeks post-MI, suggesting that these senescent cells are not entirely undergoing apoptosis and then being cleared by the immune system or then being replenished by non-senescent cells. Instead, some of them may reversibly transition to a non-senescent state, as only few p16^high^ cells (TAM administrated on days 28-30) or SA-β-Gal positive cells were observed on 4 weeks post-MI. As diverse cell types exhibit the senescent phenotype upon MI, different senescent cell types may adopt distinct cell states and survival fates during MI-induced cardiac remodeling. Further study is warranted to focus on dissecting the role of their divergent fates on cardiac function upon MI.

We systemically evaluated the composition of senescent cells in the heart following MI. Our spatial transcriptomics analysis revealed that fluorescence-labeled senescent cells were mostly localized in the ischemic region, enriched for the fibroblast marker. Then, a *Cdkn2a* score > 0.5 was used to identify senescent cells from the snMultiome RNA-seq data, demonstrating that fibroblasts and the subpopulation late MF are the largest senescent cell population. The transcription level of *Cdkn1a*, another common marker for senescent cells, was also applied to the snMultiome RNA-seq data to evaluate p21^high^ cells (*Cdkn1a* score > 0.5). Although more abundant than the p16^high^ cells, the p21^high^ cells still exhibited the consistent pattern of senescent cells (data not shown). Furthermore, by analyzing the integrated transcriptomics data, we found that all the 5 computational methods had strong consistency to conduct the cellular deconvolution task, with RCTD and Tangram exhibiting the highest overall correlation. This result may be attributed to their ability to accurately handle datasets with fine-grained cell types. In addition, Tangram is capable of performing deconvolution with large views of tissues^29^.

During days 3-7 post-MI, MF reach peak proliferation and produce significant ECM^30,31^. According to our findings, the senescent cells spiked at this time. Interestingly, not all MF became senescent. The existence of fibrogenic and non-fibrogenic, as well as other properties or subtypes within the fibroblast population, further contributes to the differential susceptibility to senescence ^32^. In our study, late MF, which appeared later in the cardiac remodeling process, are more prone to be senescent among all fibroblast subpopulations. Functional analysis indicates that senescent late MF exhibited decreased oxidative stress and lipid transport, as well as ECM production capacity. Previous studies^33,34^ have shown that cardiac fibroblasts with increased lipids uptake and utilization are required for ECM synthesis following MI through enhanced fatty acid oxidation. The absence of these functions in the senescent MF reduced ECM production capability in HCF in our *in vitro* soluble collagen assay, which may rescue the adverse effect of non-senescent late MF-induced excessive fibrosis. Furthermore, we showed that administration of senolytic drug ABT263 led to increased fibrosis and aggravated cardiac dysfunction post-MI, which indirectly supported our *in vitro* findings. In a transverse aortic constriction mouse model, increased fibrosis was observed in p53- and p16^Ink4a^-deficient mice^35^, suggesting that MF senescence may be a general essential anti-fibrotic mechanism. However, a limitation of ABT263^36^ is the lack of selectivity, resulting in elimination all types of senescent cells. This may produce a net effect, obscuring the specific impact that senescent fibroblasts have on cardiac function. Further study may focus on the effects of specifically targeting p16 positive fibroblasts on cardiac function in response to MI.

## Methods

### The myocardial infarction (MI) mouse model and senolytics administration

The experimental procedures were approved by the Institutional Animal Care and Use Committee (IACUC) at City of Hope. All mice were maintained in the central animal facility of City of Hope Beckman Research Institute. Mice used in the study were of the C57BL/6N background, aged 10-14 weeks old, and male. The temperature of mice was controlled using a heat pad (37°C) during the MI procedure. Mice were anesthetized and underwent trachea intubation. Chest opening was performed at the intercostal space of the 3^rd^ and 4^th^ ribs. MI was achieved by inducing ischemia by ligation of the left anterior descending (LAD) coronary artery using a 7-0 polypropylene suture. Sham-operated animals received the same procedure without ligation. All mice received buprenorphine for analgesic purpose and were allowed to recover in a ventilated incubator. Mice were terminated on 3 days, 7 days, 2 weeks, 3 weeks, and 4 weeks post-MI. Heart samples were harvested for future analysis. Navitoclax ABT263 (in 10% ethanol: 30% polyethylene glycol 400: 60% Phosal 50 PG) was administered to wild-type mice by oral gavage at 50 mg per kg body weight per day (mg/kg/d) for 4 or 7 consecutive days (days 4-7 or days 4-10 post-MI). Control mice were administered with an equal volume of vehicle (10% ethanol: 30% polyethylene glycol 400: 60% Phosal 50 PG) for 4 or 7 days.

### The p16 senescence reporter mouse model

The p16 senescence reporter mouse model was generated by crossing p16^Ink4a^-Cre^ERT2^ knock-in mice^13^ with the mT/mG mouse line^18^. The p16^Ink4a^-Cre^ERT2^ knock-in mice contain the first exon (Ex1α) of the endogenous p16^Ink4a^ gene substituted with a target gene cassette Cre^ERT2^. The Cre recombinase activity of Cre^ERT2^ can be controlled by tamoxifen administration to assess the long-term proliferative potential of labeled cells and determine their lifespan *in vivo*. The p16^Ink4a^-Cre^ERT2^ mice were then intercrossed with the mT/mG mouse line (#007576, Jax)^18^, resulting in p16^Ink4a^-Cre^ERT2^-mT/mG mice. The senescence reporter mouse model allowed for effective labeling of p16^high^ cells with EGFP expression specifically in the presence of 3 days of tamoxifen injection before harvest at different time points post-MI. For lineage tracing, as the p16^high^ cells presented on day 7 post-MI, TAM was administrated on days 5-7, and heart tissue was harvested 4 weeks post-MI.

### Echocardiography

Cardiac function was evaluated with unconstrained, conscious mice using echocardiography (#Vevo 3100, MS400 transducer, Visual Sonics) as previously described^37^. M-mode images of the short-axis view at the level of papillary muscles were captured and used to analyze and calculate various cardiac functional parameters. Heart rate was recorded. Left ventricular internal diameters (in mm) at end diastole (LVID, diastolic) and end systole (LVID, systolic) were determined with M-mode recordings. Ejection fraction (EF) = [(Left ventricular end-diastolic volume (LVEDV) - Left ventricular end-systolic volume (LVSV)) / LVEDV] x 100%.

### Masson trichrome staining

Cryosections of OCT heart samples (10 μm) were used for trichrome staining according to the manufacturer’s instructions (#150686, Abcam). Briefly, slides were placed in Bouin’s solution pre-heated to 56-64°C for 60 minutes and rinsed until the tissue was clear. Slides were stained with the working Weigert’s Iron Hematoxylin for 5 minutes. After washing, slides were stained with Biebrich scarlet/acid fuchsin solution for 15 minutes and then rinsed in distilled water. Slides were differentiated in phosphomolybdic/phosphotungstic acid solution for 10-15 minutes. Without rinsing, Aniline Blue solution was applied for 5-10 minutes, followed by rinsing in distilled water. Finally, slides were treated with 1% acetic acid solution for 3-5 minutes. Slides were then dehydrated in ethanol, cleared in xylene, and mounted in a resinous medium. Finally, slides were scanned with the Leica microscope, and images were analyzed with the MIQuant software^38^.

### *In vitro* fibroblast culture and treatment

Human cardiac fibroblasts (HCF) isolated from the ventricle of the adult heart were purchased from PromoCell (#C-12375) and cultured in Fibroblast Growth Medium 3 (#C-23025, PromoCell) at 37°C with 5% CO_2_. HCF were grown to confluence and growth-arrested in 1% serum medium for 24 hours before treatment with 1% serum medium alone or same medium containing 10 ng/mL TGF-β1 for 72 hours or simulated ischemia. Simulated ischemia was performed by culturing the cells in the buffer containing 117 mM NaCl, 12 mM KCl, 0.9 mM CaCl_2_-2H_2_O, 0.49 mM MgCl_2_-6H_2_O, 4 mM HEPES, 20 mM sodium lactate, and 5.6 mM 2-deoxyglucose (pH 6.2) for 6 hours in a 95% N_2_ and 5% CO_2_ chamber at 37°C and then replaced by normal culture medium (1% serum DMEM) under normoxia condition for 72 hours.

### Immunoblotting

The ischemic part of the heart tissue was lysed in ice-cold lysis buffer with proteinase inhibitors. Protein concentration was determined using the Pierce BCA Protein Assay Kit (#10741395, Thermo Fisher Scientific). Approximately 10 μg of total proteins was loaded onto an SDS-polyacrylamide gel electrophoresis gel and subsequently transferred to a nitrocellulose membrane (#162-0112, Bio-Rad Laboratories). For primary antibody incubation, the membrane was kept at 4°C overnight with one of the following antibodies: p16^Ink4a^ (1:1,000, #ab211542, Abcam), p21 (1:1,000, #2947, Cell Signaling Technology), p53 (1:20,000, #2524, Cell Signaling Technology), Phospho-Histone H2A.X (Ser139) (1:1000, #9718, Cell Signaling Technology), and GAPDH (1:1,000, #2118, Cell Signaling Technology). Secondary antibodies, including goat anti-mouse Alexa Fluor™ 680 (#A-21057, Invitrogen) and goat anti-rabbit Alexa Fluor™ 800 (#A32735, Invitrogen) diluted at 1:15,000, were applied for 1.5 hours at room temperature. Signal detection was achieved using the Li-COR Imaging system. The intensity of bands was quantified using the ImageJ software and normalized to the respective GAPDH housekeeping gene.

### Immunofluorescence staining

Hearts were embedded in OCT solution and stored in −80°C. For immunofluorescence staining, cryo-sectioned heart slice was fixed for 15 minutes with 4% paraformaldehyde (PFA) at room temperature and then permeabilized with 0.5% Triton X-100 (Thermo Fisher Scientific) for 15 minutes. Sections were washed with PBS, blocked using 2% BSA for 45 minutes, and incubated with the following primary antibodies: γH2AX (1:500, #9718S, Cell Signaling), p16^Ink4a^ (1:100, #ab211542, Abcam), α-SMA (1:100, #ab5964, Abcam), and vimentin (1:200, #sc-6246, Santa Cruz Biotechnology) at 4°C overnight. Sections were next incubated with secondary goat anti-mouse IgG or goat anti-rabbit IgG antibody (1:500, Invitrogen) for 1.5 hours at room temperature. Coverslips were then mounted using the ProLong™ Gold Antifade Mountant with DAPI (#P36935, Thermo Fisher Scientific). Images were acquired using a Leica microscope and processed using the ImageJ software.

### SA-β-Gal and SPiDER-β-Gal staining

SA-β-Gal staining was performed according to the manufacturer’s instructions (#9680, Cell Signaling Technology). Briefly, tissue slides or cells were washed in PBS buffer, fixed for 15 minutes, washed with PBS, and then incubated with fresh SA-β-Gal staining solution (pH=6) at 37°C without CO_2_ overnight. For SPiDER-β-Gal staining, fixed frozen heart tissue slices used for spatial transcriptome analysis were directly incubated with 20 μmol/L SPiDER-β-Gal reagent (#SG02, Dojindo) at room temperature for 30 minutes. The OCT-embedded fresh frozen heart tissue used for regular SPiDER-β-Gal staining was fixed with 4% PFA for 15 mins and then incubated in the SPiDER-β-Gal reagent at room temperature for 30 min. After incubation, the samples were washed with PBS, observed under a Leica microscope, and processed with the ImageJ software.

### Visium tissue preparation and sequencing

The 10x Genomics Visium Spatial Gene Expression platform enables the integration of immunofluorescence staining with spatial transcriptomics analysis in cryo-sectioned tissue slices, allowing for the identification of transcriptomic profiles in staining-positive cells. We applied this technique to the fixed, frozen heart slices from wild-type mice, using SPiDER-β-Gal staining to label senescent cells. SPiDER-β-Gal staining relies on an enzymatic reaction to conjugate fluorescent compounds, different from the conventional immunofluorescence staining which requires two consecutive days. We reduced SPiDER-β-Gal staining time to 30 minutes to preserve the morphological quality of tissue sections and the integrity of mRNA transcripts. Briefly, the heart was divided into two parts below the ligature. The upper part of the heart tissue was fixed and flash-frozen in an isopentane bath before embedding in OCT, which was then sectioned at 10 μm and placed on 10x Visium gene expression slides (11 mm x 11 mm capture area) according to the 10x Visium CytAssist Spatial Gene Expression for Fixed Frozen-Tissue Preparation, Imaging & Decrosslinking Guide to perform SPiDER-β-Gal staining. Similarly, the infarct slice labeled with EGFP from the upper part of the heart on day 7 post-MI of the p16 senescence reporter mouse was used for spatial profiling. cDNA libraries with spatial tagging were built using the Visium Spatial Gene Expression 3’ Library Construction v1 Kit (10x Genomics) and sequenced using a NovaSeq 6000 sequencing system (Novogene).

### Visium data processing and quality control

Manual fiducial alignment and tissue outlining were performed using Loupe Browser (v6.1.0). Samples were processed with SpaceRanger (1.3.1) based on mouse reference genome mm10 (reference package refdata-gex-mm10-2020-A). SpaceRanger was also used to align paired histology images with mRNA capture spot positions in the Visium slides. SpaceRanger output was further processed with Seurat package^39^, which includes normalization to preprocess raw expression data, dimensional reduction, and clustering to identify distinct cell populations and spatially-variable features to highlight genes with spatial patterns, as well as interactive visualization by utilizingdplyr, ggplot2, and patchwork packages under R (v4.2.3).

### Single-nucleus Multiome (snMultiome) sequencing

Flash-frozen heart samples from wild-type and p16 senescence reporter mice on day 7 post-MI were used for snMultiome sequencing. The heart was cut below the ligature, and the lower part of the heart (left ventricle, excluding the septum) was collected and triturated. snMultiome-seq libraries were prepared using the 10x Genomics Single Cell Multiome ATAC + Gene Expression Kit according to the manufacturer’s instructions. Briefly, nuclei were isolated, and GEMs were generated by the Next GEM Chip J Single Cell Kit (10x Genomics). The quality of libraries was estimated by the 2100 Bioanalyzer Instrument (Agilent). The libraries were sequenced on an NovaSeq6000 sequencing system (Novogene). The FASTQ sequencing reads were processed and aligned to the mouse genome (mm10).

### snMultiome data processing and quality control

snMultiome ATAC and RNA sequencing fastq files were converted to digital gene expression matrices using 10x Genomics Cell Ranger ARC v2.0.2 pipeline according to the manufacturer’s recommendations (https://support.10xgenomics.com/single-cell-gene-expression/software/overview/welcome) with the mouse reference genome mm10. To eliminate batch difference and sequencing depth difference and ensure sample comparability, we pooled the results of each sample individually from cellranger-arc count and performed aggregation with cellranger-arc aggr. To conduct quality assessment of library, both RNA and ATAC data from Cellranger-arc aggr output were configured to a Seurat object using Seurat 5.0.0 and Signac 1.11.0 in R (v4.2.3). For ATAC data, peaks were called using MACS2, and the genomic positions were mapped and annotated with reference mouse genome EnsDb.Mmusculus.v79_2.99.0 and mm10. Low-quality cells, including multiplets, dead cells, and those with poor sequencing quality, were removed based on the following criteria: nCount_RNA < 1,000 or > 100,000; nCount_ATAC < 1,000 or > 150,000; nucleosome signal > 2; or TSS enrichment < 2. Further filtering of doublets was performed with doubletFinder. Gene expression data were processed with SCT transform. After filtering low quality cells, we removed peaks on nonstandard chromosomes and in genomic blacklist regions from ENCODE.

### Cell clustering and cell type annotation

Gene expression (RNA) data were normalized using SCTransform while regressing out sample-specific effects. The top 50 principal components (PCs) were extracted and used to compute the 30 nearest neighbors based on cosine distance. This neighbor graph was subsequently used for UMAP dimensionality reduction with default parameters. For the ATAC data, term frequency–inverse document frequency (TF-IDF) normalization was applied, followed by selection of the top features and dimensionality reduction using singular value decomposition (SVD). UMAP was then performed using the resulting components, excluding the first dimension which primarily captures sequencing depth. Next, we calculated a weighted-nearest neighbor (WNN) graph, representing a weighted combination of RNA and ATAC-seq modalities. FindMultimodalNeighbors with gene expression based pca and ATAC based lsi reductions were used to generate a joint neighbor WNN graph. FindClusters with SLM algorithm was used to identify clusters. Cell type annotation was manually performed by identifying cluster-specific biomarkers against all other clusters with FindMarkers function. The specific markers are listed as below: cardiomyocytes (*Ryr2, Tnnt2, Actn2*), endothelial cells (*Vegfc, Tie1, Pecam1*); fibroblasts (*Postn, Dcn, Col1a1*), macrophages (*C1qb, C1qa, Cd68*), epicardial cells (*Bnc1, Wt1, Msln*); smooth muscle cells (*Acta2, Tagln, Myh11*); pericytes (*Kcnj8, Rgs5, Abcc9*), T cells (*Cd3g, Cd8a, Cd3e*), B cells (*Cd79a, Cr2, Igkc*), and dendritic cells (*Flt3, Ccr7, Xcr1*). For fibroblasts subpopulations, the following markers were used: uninjured quiCF (*Hmcn2, Col6a6*), acute quiCF (*Mt2, Ror1*), early ActCF (*Lars2, Camk1d*), acute ActCF (*Csrp1, Spp1*), early MF (*Top2a, Neil3*), late MF (*Cacna1c, Postn*), and MaF (*Itgbl1, Nox4*), and chronic quiCF (*Col8a1, Prkca*).

### Differential peak and gene expression analysis

Differential peak and gene expression analysis was performed based on the non-parametric Wilcoxon rank sum test with FindMarkers function between groups. Upregulated genes and peaks in senescent cells versus non-senescent cells were filtered with avg_log2 folder change > 0.5 and adjusted *p* < 0.05. Downregulated genes and peaks were filtered with avg_log2 folder change < −0.5 and adjusted *p* < 0.05.

### Pathway enrichment and transcription factor (TF) motif enrichment analysis

To explore the functional implications of the senescent cells in MI, we first performed pathway enrichment analysis using clusterProfiler (version 4.6.2). Next, we explored the key transcription factor (TF) regulating senescence by examining the accessible regions of each cell state to determine enriched motifs in senescent cells. As described in the Signac motifs vignette, we called peaks using MACS2 and linked them to nearby genes by assessing the correlation between peak signals and gene expression. For each gene, this function computed the correlation coefficient between gene expression and accessibility of each peak within a given distance from transcription start site. It also computed the expected correlation coefficient for each peak based on its GC content, accessibility, and length. The expected coefficient values for the peak were then used to compute a z-score and p-value. RunChromVAR of chromVAR 1.12.0 package and JASPAR version 2020 database were used to compute motif enrichment. Promoter accessibility and motif enrichment information were incorporated as the assay data in the integrated Seurat object files.

### Projection of spatial transcriptomics dataset to snMultiome dataset

To project the SPiDER-β-Gal/p16-EGFP fluorescence positivity from spatial transcriptomics dataset to snMultiome-seq dataset, we first identified the fluorescence positive or negative spots from the wild-type mouse and p16 senescence reporter mouse heart on day 7 post-MI and generated two fluorescence status annotations for spatial transcriptomics datasets.

Subsequently, Seurat function FindVariableFeatures was used to identify the molecular features that were specific for each spot in spatial transcriptomics dataset as well as specific for each nucleus in snMultiome dataset. By using the shared variable features from both datasets, we applied the anchor-based integration workflow in Seurat, which enabled the probabilistic transfer of annotations (fluorescence status) from the reference (spatial transcriptomics) to the query (snMultiome-seq) datasets.

### Implementation of 5 algorisms

To identify the cell types of senescent cells labeled with SPiDER-β-Gal/p16 EGFP fluorescence, we applied 5 algorithms to deconvolute the cellular composition within each spot from the Visium spatial transcriptomics dataset using the snMultiome RNA-seq dataset as reference.

Cell2location: Version 0.1.4 was used following the tutorial on https://cell2location.readthedocs.io/en/latest/notebooks/cell2location_tutorial.html. We used N_cells_per_location=10 and detection_alpha=20.

Tangram: Version 1.0.4 was used following the tutorial on https://github.com/broadinstitute/Tangram/blob/master/tutorial_tangram_with_squidpy.ipynb. We used mode=“cells”.

SPOTlight: Version 1.10.0 from Bioconductor was used following the tutorial on https://www.bioconductor.org/packages/release/bioc/vignettes/SPOTlight/inst/doc/SPOTlight_ki dney.html. We used n_cells=100.

RCTD: Spacexr 2.2.1 (which includes implementations of RCTD and C-SIDE) from Bioconductor was used following the tutorial on https://raw.githack.com/dmcable/spacexr/master/vignettes/spatial-transcriptomics.html. We used doublet_mode=“full”, which means that no restriction is placed on the number of cell types per spot.

CellDART: Version 0.1.3 was used following the tutorial on https://github.com/mexchy1000/CellDART.

### Statistical analysis

Data presented in this study were representative of at least 3 independent experiments and expressed as mean ± standard deviation (SD). Kolmogorov-Smirnov test was used to test if the data is normally distributed. A *p* >0.05 on test indicated that the data are viewed as normally distributed. For normally distributed data, differences between two experimental groups were assessed using unpaired, two-tailed Student’s t-test. Differences between three or more groups were tested by one-way or two-way ANOVA followed by Tukey post-hoc test for inter-group comparison. Alternatively, a two-tailed Wilcoxon rank-sum test for two groups and two-tailed Kruskal–Wallis test for three or more groups were used when data were not normally distributed. The agreement between each pair of deconvolution algorithms was analyzed using Pearson correlation coefficient. The group size for each experiment is provided as ‘n’. All replicates are biological replicates. A statistically significant difference was defined as *p* < 0.05. Except for snMultiome data, all data analysis were implemented in GraphPad Prism 9.0 software.

## Data availability

The snMultiome RNA-sequencing and ATAC-sequencing and spatial transcriptomic data are available through the Gene Expression Omnibus database (GSE296300 and GSE296301). All software scripts are published in GitHub: https://github.com/ZhaoWangLab/Jielin-Deng_Multi-omics-in-MI.git.

### Please note

*for reviewing purpose, the snMultiome dataset was uploaded to NCBI with GEO296300, the link to access this dataset is* https://www.ncbi.nlm.nih.gov/geo/query/acc.cgi?acc=GSE296300, *and the token to enter for access is kbepuciqntuvzqr. The GEO information for spatial transcriptomics dataset is GEO296301, the link to access is* https://www.ncbi.nlm.nih.gov/geo/query/acc.cgi?acc=GSE296301, *and the token to enter is uvapmykgjlgzvap*.

## Acknowledgements

This work was supported by funding from the American Heart Association (23TPA1077069 to Z.V.W.), the American Diabetes Association (7-20-IBS-218 to Z.V.W. and 1-19-JDF-082 to Y.D.), and the National Institute of Health (R01 HL137723, R01 HL156951, and R01 HL171309 to Z.V.W. and R01 DK-126975 to Y.D.). Z.V.W. is an Established Investigator of the American Heart Association. Schematic diagram figures were prepared with biorender.com.

## Author contributions

J.D. and Z.V.W. conceived and designed the study. J.D., D.R., H.Z., Q.X., Y.J., Y.Z., Y.W., and H.W. performed experiments and data analysis. J.D., D.R., H.Z., and Y.J. interpreted the data. M.N. provided key reagents. J.D., Y.D, and Z.V.W. edited and revised the manuscript.

