## supplementary materials for "Multi-omic Profiling of Senescence Following Myocardial Infarction Reveals Dynamic Characteristics in Cardiac Remodeling"

**Letter**

Department of Diabetes and Cancer Metabolism, Fox North Room 206, Beckman Research Institute of the City of Hope, 1500 East Duarte Road, Duarte, California 91010, USA.

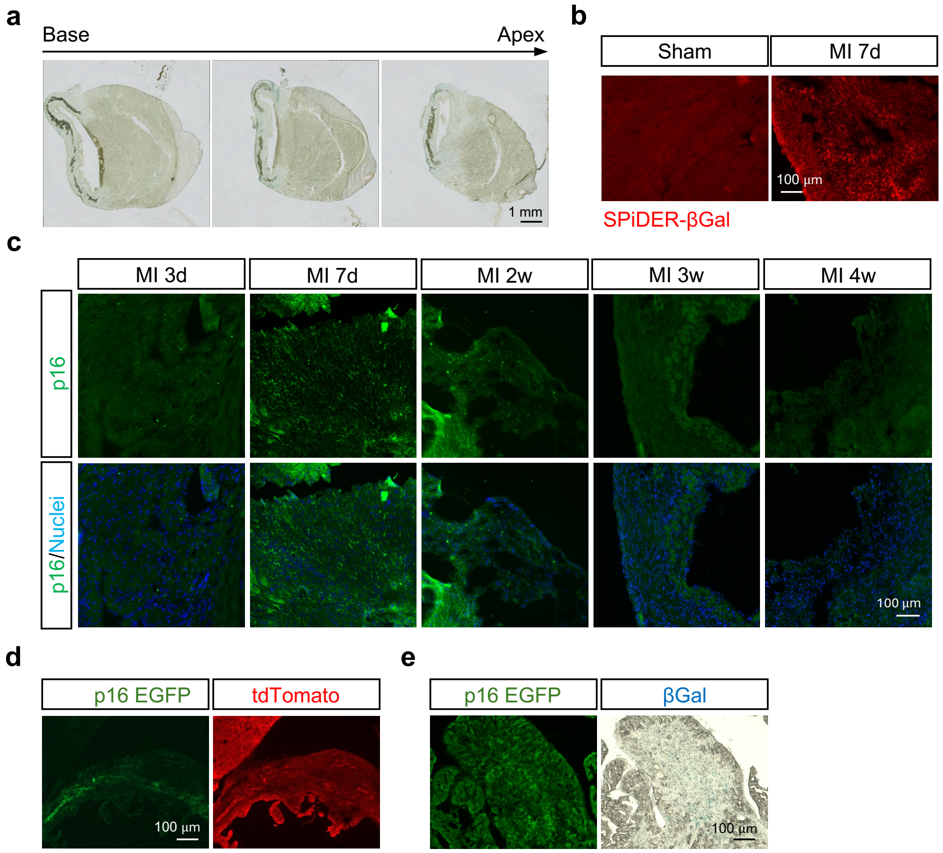


**Extended Data Figure 1. Senescence dynamics in the heart following MI.**

- **a.** Senescence-associated β-galactosidase (SA-β-Gal) staining of different heart cross-sections on day 7 post-MI, demonstrating consistent levels of senescence across multiple layers of the infarcted myocardium.
- **b.** SPiDER-β-Gal staining of heart sections showed increased activity in the infarcted region of day 7 post-MI compared to sham in wild-type mice.
- **c.** $p16^{Ink4a}$ immunofluorescence staining (green) and DAPI nuclear staining (blue) of the infarcted myocardium at different time points post-MI, highlighting increased senescence in the injured cardiac tissue.
- **d.** Whole-mount fluorescence (EGFP and tdTomato) imaging from the heart sections of the p16 senescence reporter mice on day 7 post-MI.
- **e.** Colocalization of p16-EGFP positive cells with SA-β-Gal positive cells from the same heart section of the p16 senescence reporter mice on day 7 post-MI, indicating high similarity between p16 expression and SA-β-Gal activity as the senescent markers.

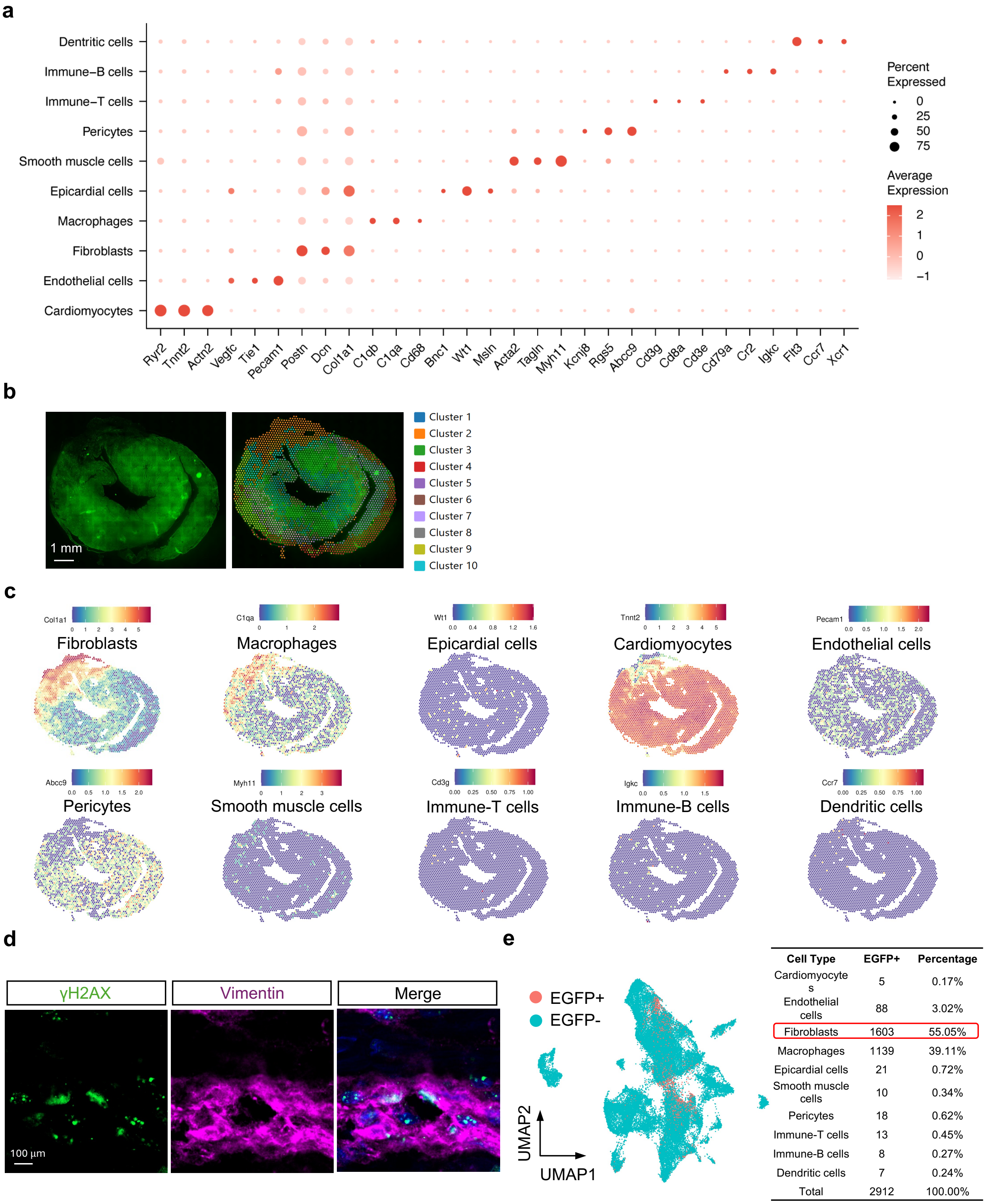


| Cell Type | EGFP+ | Percentage |
|---|---|---|
| Cardiomyocytes | 5 | 0.17% |
| Endothelial cells | 88 | 3.02% |
| Fibroblasts | 1603 | 55.05% |
| Macrophages | 1139 | 39.11% |
| Epicardial cells | 21 | 0.72% |
| Smooth muscle cells | 10 | 0.34% |
| Pericytes | 18 | 0.62% |
| Immune-T cells | 13 | 0.45% |
| Immune-B cells | 8 | 0.27% |
| Dendritic cells | 7 | 0.24% |
| Total | 2912 | 100.00% |

**Extended Data Figure 2. Fibroblasts constitute a major portion of senescent cells in the infarcted heart post MI.**

**a.** Single-nucleus (sn) Multiome sequencing analysis from infarcted hearts on day 7 post-MI in wild-type mice and p16 senescence reporter mice. WNN identified 10 cardiac cell types. Dot plot shows the expression of marker genes of each cell type. The marker genes used for identifying cell types including the follows: fibroblasts (Fb, n=18,689, 41.93%), macrophages (n=10,181, 22.84%), endothelial cells (EC, n=8,759, 19.65%), pericytes (n=2,703, 6.06%), cardiomyocytes (CM, n=1,931, 4.33%), epicardial cells (n=980, 2.2%), smooth muscle cells (SMC, n=355, 0.80%), immune-T cells (n=430, 0.96%), immune B cells (n=282, 0.63%), and dendritic cells (n=263, 0.59%).

**b.** Unsupervised spatial transcriptomics analysis of the p16 senescence reporter mouse heart on day 7 post-MI.

**c.** Fibroblasts were predominantly localized in the ischemic region (cluster 2) where the $p16^{high}$ EGFP positive senescent cells mainly located.

**d.** Immunofluorescence validation of senescent fibroblasts in wild-type mouse hearts on day 7 post-MI. γH2AX (green) colocalizes with vimentin (magenta), confirming that fibroblasts are a major senescent cell type in the heart.

**e.** Mapping the p16 EGFP fluorescence-based spatial transcriptomics to snMultiome data, showing that fibroblasts comprise 55.05% of the total SPiDER-βGal positive cells.

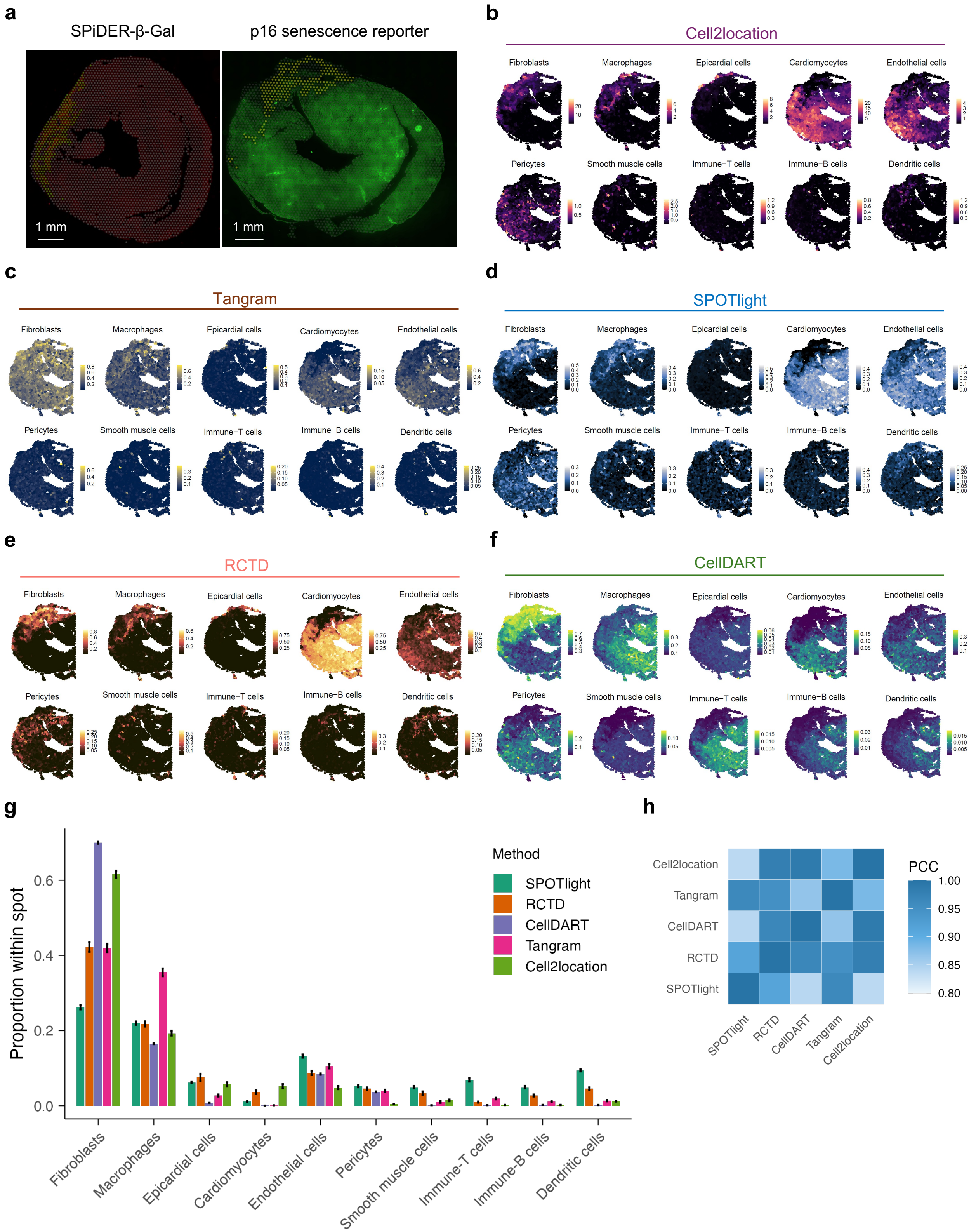

a
SPiDER-β-Gal
p16 senescence reporter
1 mm
b
Cell2location
c
Tangram
d
SPOTlight
e
RCTD
f
CellDART
Fibroblasts
Macrophages
Epicardial cells
Cardiomyocytes
Endothelial cells
Pericytes
Smooth muscle cells
Immune−T cells
Immune−B cells
Dendritic cells
g
Proportion within spot
Method
SPOTlight
RCTD
CellDART
Tangram
Cell2location
h
PCC

**Extended Data Figure 3. Cellular deconvolution of the p16 EGFP positive cells confirms fibroblasts as the predominant senescent cell type in the heart post-MI.**

**a.** A total of 254 SPiDER-β-Gal positive spots and 200 EGFP positive spots were selected manually from heart sections of the wild-type mouse and p16 senescence reporter mouse on day 7 post-MI, respectively, visualized on Visium spatial transcriptomics slides.

**b-f.** Cellular deconvolution using Cell2location (**b**), Tangram (**c**), SPOTlight (**d**), RCTD (**e**) and CellDART (**f**) to estimate the proportion of the Visium spots in the infarcted heart from the p16 senescence reporter mouse on day 7 post-MI.

**g.** Statistical analysis of cell type compositions predicted by the 5 deconvolution algorithms in EGFP positive spots confirmed fibroblasts as the predominant senescent cell type.

**h.** Pearson correlation coefficients (PCC) were calculated to assess the agreement between each pair of the algorithms in predicting EGFP positive senescent cell types in the infarcted p16 senescence reporter mouse heart. Among the 5 algorithms, RCTD showed the highest overall correlation with the others. Average PCC values were as follows: SPOTlight, 0.89; RCTD, 0.95; CellDART, 0.92; Tangram, 0.92; Cell2location, 0.92.

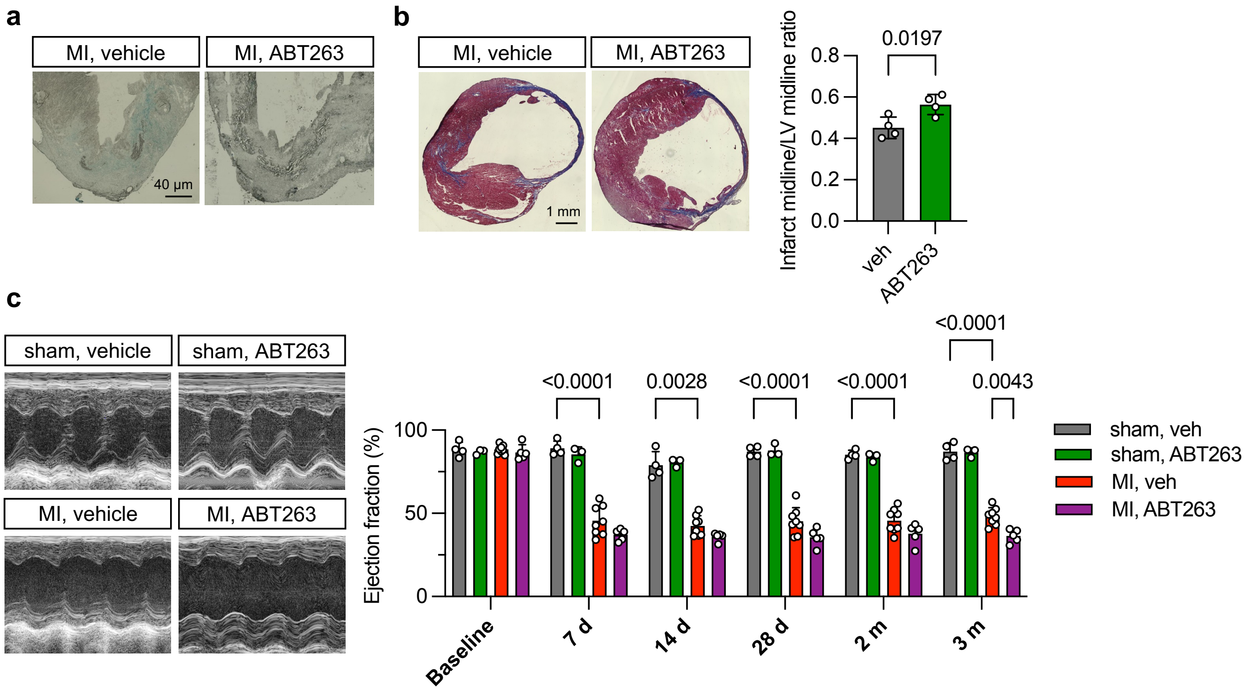


**Extended Data Figure 4. Depletion of senescent cells from the heart exacerbates cardiac dysfunction post-MI.**

- **a.** ABT263 administration (days 4-7 post-MI) significantly reduced the SA-β-Gal positive cells in the heart compared to the vehicle-treated group on day 7 post-MI.
- **b.** ABT263 administration (days 4-10 post-MI) increased infarct size and collagen deposition (Masson's trichrome staining) in wild-type mice 3 months post-MI compared to the vehicle-treated mice, as indicated by midline length (calculated by dividing the midline length of the infarcted left ventricular wall by the midline length of total left ventricular wall). MI + vehicle, n=4; MI + ABT263, n=4. Unpaired t-test was conducted. Data are represented as mean ± SD.
- **c.** Representative cardiac echocardiographic images are shown. ABT263 administration (days 4-10 post-MI) caused a decrease in ejection fraction 3 months post-MI in the wild-type mice, as measured from echocardiographic recordings. sham + vehicle, n=4; sham + ABT263, n=3; MI + vehicle, n=8; MI + ABT263, n=5. Two-way ANOVA followed by Tukey post-hoc multiple comparisons test was conducted. Data are represented as mean ± SD.
